# Persistent cerebrovascular injury after preeclampsia in the STOX1A mouse model: implications for prokineticin signaling

**DOI:** 10.64898/2026.10.29.762861

**Authors:** Hadi Younes, Frederic Sergent, Ioanna Kyritsi, Lise Miguel, Olivier Montigon, Daniel Vaiman, Mohamed Benharougha, Emmanuel Barbier, Nadia Alfaidy, Christel Marquette

## Abstract

Preeclampsia (PE) is a pregnancy-related hypertensive disorder associated with neurological complications that may persist beyond pregnancy. Prokineticins (PROK1 and PROK2) and their receptors (PROKR1 and PROKR2) regulate angiogenesis and inflammation and may be involved to PE-associated cerebrovascular injury. This study investigated alterations of the maternal brain prokineticin system during PE and 8 months postpartum, and assessed the consequences of prokineticin receptor inhibition using the PROKR antagonist PC1. Female WT FVB/N mice mated with transgenic males overexpressing STOX1A were used as a model of PE and received PC1 or vehicle during gestation. Circulating and regional brain PROK levels and receptor expression were assessed by ELISA and Western blotting. Long-term cerebral abnormalities were evaluated using high-resolution MRI and GFAP immunostaining. PE mice exhibited hypertension, proteinuria, reduced placental efficiency, and elevated circulating PROK1 and PROK2 levels. PC1 prevented hypertension and attenuated the elevation of circulating PROKs but did not prevent proteinuria or reduced placental efficiency. During PE, hippocampal PROK1 increased while PROK2 decreased; these changes persisted despite treatment. At 7 months postpartum, both PROKs were elevated in the cortex, and these increases were prevented by PC1. Region-specific changes in PROKR1 immunoreactivity and PROKR2 expression showed variable responses to treatment. MRI revealed hippocampal and parieto-temporal T2* hypointensities compatible with microhemorrhagic injury, post-contrast T1 abnormalities supporting blood-brain barrier disruption, and a distinct caudoputamen T1-hyperintense lesion potentially representing a nonhemorrhagic component. PC1 prevented focal cerebral lesions and associated astrocyte reactivity, whereas regional diffusion abnormalities persisted. These findings implicate prokineticin signaling in persistent PE-associated brain injury and support the potential of receptor antagonism to limit selected cerebral alterations.

## 1. Introduction

Preeclampsia (PE) is a multisystem hypertensive disorder of pregnancy characterized by the new-onset of hypertension (systolic blood pressure ≥ 140 mmHg and/or diastolic blood pressure ≥ 90 mmHg) after 20 weeks of gestation, often accompanied by proteinuria (spot urine protein/creatinine ratio ≥ 30 mg/mmol, ≥ 300 mg/24 h, or ≥ 1 g/L [2+] on dipstick testing) or evidence of maternal or uteroplacental dysfunction [1, 2]. The International Society for the Study of Hypertension in Pregnancy (ISSHP) has refined these diagnostic criteria to improve clinical practice and research standardization, emphasizing the heterogeneous nature of PE and its multifactorial pathophysiology [1]. Despite advancements in the management of PE, it remains a leading cause of maternal and perinatal morbidity and mortality worldwide [3].

Beyond its acute effects, PE is increasingly recognized as a risk factor for long-term neurological complications particularly cognitive decline and dementia. Cohort studies associate a history of PE with an increased risk of later-life dementia, particularly vascular dementia. Cognitive studies also report difficulties in memory, executive function, and processing speed, although objective findings vary across studies [4–10]. Additionally, neuroimaging studies have revealed white matter lesions (WML) in the frontal and temporal lobes of women with a history of PE, which may reflect vascular injury [11–13]. These findings suggest that PE may induce persistent cerebral alterations that contribute to long-term cognitive deficits.

The molecular and cellular mechanisms underlying these neurological sequelae remain poorly understood. However, emerging evidence implicates neuroinflammation, blood-brain barrier (BBB) dysfunction and dysregulation of angiogenic and inflammatory mediators in neurological manifestation of PE [14–19]. The BBB, a critical interface maintaining cerebral homeostasis, is compromised in PE as evidenced by alterations in tight-junction proteins, increased permeability and neuroinflammatory responses [14–17]. These changes can contribute to cerebral edema and vasogenic injury and may be involved in long-term neurological impairments.

Among the emerging mediators in PE, the prokineticin family, particularly Prokineticin 1 (PROK1, also known as Endocrine Gland-Derived Vascular Endothelial Growth Factor, EG-VEGF), has been identified as highly dysregulated in this condition [20, 21]. The prokineticin system comprising PROK1, PROK2 and their G-protein-coupled receptors (PROKR1 and PROKR2) plays a pleiotropic role in angiogenesis, immune modulation, and reproductive physiology [22, 23]. Beyond reproduction, the prokineticin system is implicated in neurogenesis, circadian rhythm regulation, pain modulation, and neurodegenerative diseases (e.g., Alzheimer’s and Parkinson’s) [24–31]. Given its dual role in angiogenesis and inflammation the prokineticin system represents a potential therapeutic target for conditions characterized by vascular dysfunction and neuroinflammation, including PE.

Previous experimental studies have demonstrated persistent cerebral inflammation, vascular dysfunction, and memory impairment after PE in rats [19, 32]. Clayton et al. induced placental ischemia by surgically reducing uterine perfusion pressure, whereas Johnson et al. induced a PE-like phenotype using a high-cholesterol diet. Nevertheless, these models induced the disease through surgical or dietary interventions rather than a primary genetic perturbation of the trophoblast function. In this study, we used the well-established mouse model of PE, the STOX1A-PE mouse model that was initiated by fetoplacental overexpression of human STOX1A, a transcription factor directly implicated in trophoblast differentiation and invasion [33, 34]. STOX1A regulates trophoblast migration and invasion as well as managements of oxidative/nitrosative stress in the placenta [34, 35]. In human trophoblast models, the PE-associated Y153H variant reduces invasion through upregulation of the cell-cell adhesion protein alpha-T-catenin (CTNNA3) [36]. Inadequate trophoblast invasion can impair spiral artery remodeling and uteroplacental perfusion, thereby promoting placental hypoxia [2]. In the TgSTOX42 model, the severe PE phenotype [33] is associated with a hypoxic placental environment. Using this genetic model, we studied the long-term neurological consequences of a hypertensive pregnancy, induced by the STOX1A model, focusing on brain damage, neuroinflammation associated with astrogliosis, and prokinetin dysregulation. We examined the levels of expression of PROK1 and PROK2 in blood and brain tissue both during the preeclamptic crisis and 8 months postpartum and assessed brain lesions using MRI. Additionally, we tested the preventive effects of PC1, a PROKR antagonist in attenuating PE-associated neurological adverse effects

Our findings demonstrate that PE in the STOX1A mouse model is associated with persistent cerebral abnormalities including MRI features compatible with hemorrhagic/non-hemorrhagic lesions and microstructural changes, BBB permeability alteration and neuroinflammation as evidenced by GFAP immunostaining. Furthermore, gestational PC1 treatment effectively reduced PE-associated brain abnormalities highlighting its potential for mitigating the long-term neurological sequelae of PE. These findings provide new insights into mechanisms linking PE with persistent cerebrovascular alterations and identify the prokineticin system as a candidate for further therapeutic investigations.

## 2. Materials and Methods

### Animal model

All animal experiments were conducted in strict accordance with the institutional guidelines and the European Community regulations for the use of experimental animals (Ministerial Agreement APAFIS #41782-2023031713542907_v1). Female FVB/N mice (2–4 months old. Charles River, Saint Quentin Fallavier, France) were mated with transgenic male STOX1AA mice of the TgSTOX42 line (FVB/N background, kindly provided by Dr. Vaiman’s laboratory, Cochin Hospital, France) to reproduce severe preeclampsia (PE) symptoms [33]. Control pregnancies were generated by mating FVB/N females with FVB/N males.

Throughout gestation, mice were housed in a controlled environment with a 12-hour light/dark cycle constant temperature and ad libitum access to food and water. Body weights were recorded daily during gestation. At 18.5 days of gestation (E18.5) placentas and fetuses were weighed, and the fetal-to-placental weight ratio (an indicator of placental efficiency) was calculated and compared across groups (PE, PE-PC1, and control). To confirm hypertensive gestation urinary albumin-to-creatinine ratios were assessed at E7.5 and E14.5 using metabolic cages. Urine samples were collected over 12-hour periods. Additionally, noninvasive tail-cuff blood pressure measurements were obtained daily from plug detection to E18.5 using the CODA High Throughput System (Kent Scientific, Emka Technologies, France).

### Treatment with prokineticin receptor antagonist (PC1)

To determine the role of prokineticins in PE pathology, mice were treated with PC1, a dual antagonist of prokineticin receptors 1 and 2 (kindly provided by Dr. Balboni [37]). Three experimental groups were established: the preeclampsia group injected with PC1 vehicle (1% DMSO in NaCl), the treated preeclampsia group receiving 10 intraperitoneal injections of PC1 (300 µg/kg) from E7.5 to E18.5, and the control group constituted with FVB/N females mated with FVB/N males and injected with PC1 vehicle. Two independent experiments were performed, each including the three groups. In the first experiment, mice were sacrificed at E18.5 (n = 6 per group). In the second experiment, mice were allowed to age until 8 months postpartum (n = 6 per group).

### Measurements of Prokineticin 1 and 2 concentrations

Mice were anesthetized with ketamine (100 mg/kg) and xylazine (10 mg/kg) prior to intracardiac blood collection in EDTA-coated tubes. Brains were rapidly dissected on ice into regions: cortex, hippocampus, thalamus, hypothalamus, and cerebellum, and stored at -80°C until analysis.

Protein extraction was performed using a lysis buffer (50 mM Tris-HCl pH 7.5, 150 mM NaCl, 1% Triton X-100, 0.1% SDS). After centrifugation, protein concentrations were determined using the Micro BCA Protein Assay Kit (Thermo Scientific, #23235). PROK1 and PROK2 concentrations in brain tissues and plasma were quantified using ELISA kits (Biorbyt, #orb776343 for PROK1; #orb56537 for PROK2). The intra- and inter-assay coefficients of variation (CV) were < 10% with detection limits of 15.63 pg/mL (PROK1) and 0.313 ng/mL (PROK2). For brain tissues, 300 µg of protein were used, while plasma samples (100 µL) were analyzed undiluted. Absorbance at 450 nm was measured using a TECAN Spark ELISA reader.

### Prokineticin receptor 1 and 2 expression analysis

Western blotting was performed to assess PROKR1 and PROKR2 expression in brain tissues. Thirty micrograms of protein were loaded onto 4–20% precast polyacrylamide gels (Mini-PROTEAN TGX Stain-Free™, Bio-Rad) and transferred into 0.2 µm nitrocellulose membranes (Bio-Rad, #1704271). Membranes were blocked with EveryBlot Blocking Buffer (Bio-Rad, #12010020) for 10 minutes and incubated overnight at 4°C with primary antibodies: PROKR1 (1:200, 0.2 µg/mL, Covalab, Lyon, France), PROKR2 (1:200, 0.53 µg/mL, Covalab, Lyon, France), and β-actin (1:1000, clone AC-15, Sigma-Aldrich, #A5441) as a loading control. After washing, membranes were incubated with HRP-conjugated secondary antibodies (1:1000, Jackson ImmunoResearch, #115-035-062 for anti-rabbit, #111-035-144 for anti-mouse) for 1 hour at room temperature. Chemiluminescent signals were detected using the ECL Western Blotting Substrate (Bio-Rad, #1705061) and captured with an iBright FL1000 imager (Invitrogen). β-actin was used as a standard for protein loading, and normalization was performed using ImageJ 1.53a software.

### Magnetic resonance imaging acquisitions

Eight months postpartum, anatomical MRI acquisitions were carried out at 9.4 T (Biospec Avance III HD, Bruker, Ettlingen, Germany; IRMaGe facility) using a receive-only cryoprobe dedicated to imaging the mouse brain. Animal experiments were conducted in strict accordance with the regulations and guidelines of the European Community for the use of experimental animals (Ministerial Agreement APAFIS #41782-2023031713542907_v1). Animals were anesthetized with isoflurane at approximately 1.5% in oxygen-enriched air (30% O₂) to prevent artifacts on the acquired images due to animal movement. A catheter was placed in the tail vein to allow contrast agent injection during MRI acquisition. The mice were positioned prone on a specific cradle adapted to the diameter of the magnet. A heating blanket was placed under the animals to keep them warm throughout imaging, and a sensor for recording the respiratory frequency was positioned under their thorax. The cradle was then inserted into the magnet for MRI acquisition.

The following images were collected: T2-weighted MRI to detect lesions such as edema, T2*-weighted MRI to detect bleeds, 3D T1-weighted gradient-echo MRI, and diffusion-weighted imaging to detect lesions such as edema. Three minutes after injection of Gd-DOTA (200 µmol/kg, 100 µL) (Clariscan, GE Healthcare), the 3D T1-weighted imaging sequence was repeated to evaluate blood-brain barrier integrity. Regarding data processing, diffusion-weighted data were processed using MRtrix (v3.0_RC3-137 [38]). After image denoising, unringing, and brain masking, tensor-derived fractional anisotropy and mean diffusivity maps were generated. For the determination of vascular wall integrity, contrast enhancement was calculated by subtracting the T1-weighted image before injection from the T1-weighted image obtained after injection.

### Immunohistochemistry

Immunohistochemistry for glial fibrillary acidic protein staining was performed. Fixed brains were sectioned with a vibratome and mounted on polylysine adhesion microscope slides. Brain sections were pretreated with 1% MetOH-H₂O₂ for 20 minutes at room temperature, incubated for 2 hours at room temperature with anti-GFAP antibody (1/1000, InVitrogen), and visualized with 3,3-diaminobenzidine tetrahydrochloride by the avidin–biotin–peroxidase complex method (Vectastain Elite ABC peroxidase kit, Vector Laboratories). Tissues were counterstained with hematoxylin and eosin and imaged under optical Zeiss Axioplan microscope. The acquired images were analyzed using ImageJ software.

### Statistical analysis

Data analysis of ELISA tests and Western blots were conducted using GraphPad Prism version 11.0.2. Multiple comparisons were statistically evaluated using one-way or two-way ANOVA with Tukey or Sidak post-tests, with p-values less than 0.05 considered significant. MRI statistical analyses were performed on the “eroded” data and are expressed as mean ± standard error of the mean using GraphPad Prism version 11.0.2. Mann-Whitney tests were performed to determine statistical significance, with a p-value < 0.05 considered significant.

## 3. Results

### Preeclamptic mice and PC1 treatment

According to our experimental design, mice were subjected to various assessments throughout the gestation period to validate that they underwent a hypertensive gestation (Figure 1A). To decipher a potential therapeutic approach to prevent preeclampsia, a specific treatment with PC1, an antagonist of the prokineticin receptors 1 and 2 was used, by injection between day 7.5 and day 18.5 during preeclamptic gestation.

**Figure 1:**
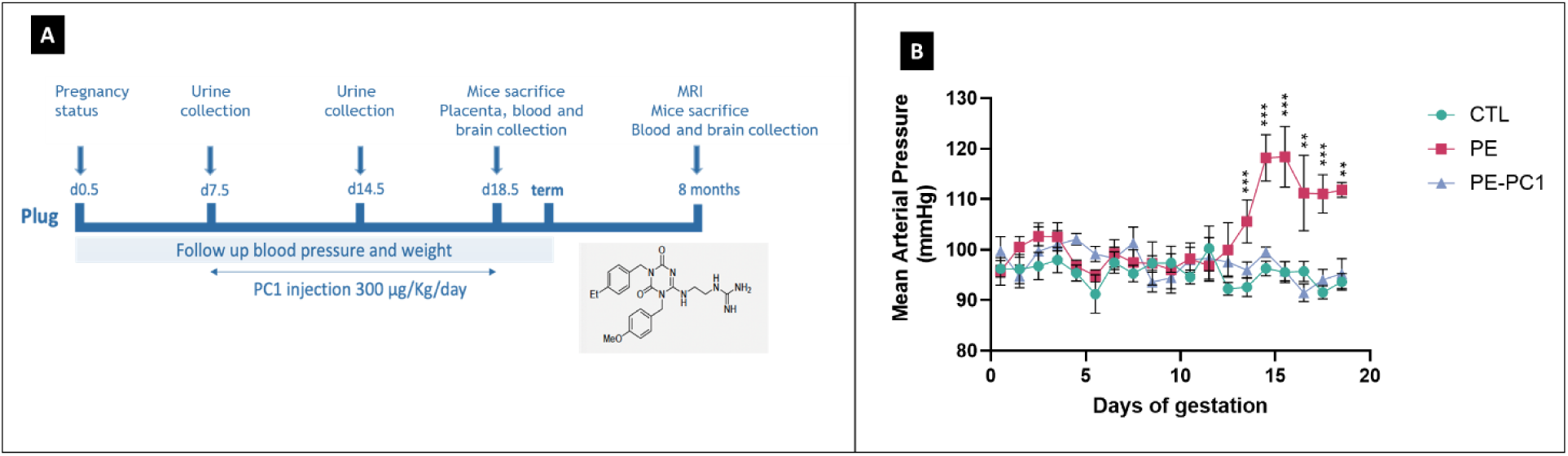
Experimental design of preeclamptic treatment with the prokineticin receptor antagonist during gestation and 8 months postpartum follow-up. (A) Experimental procedures included blood pressure and weight monitoring, with urine samples collected at 7.5 and 14.5 days post-coitum (dpc). Treated mice received PC1 via intraperitoneal injection from 7.5 to 18.5 dpc. At 18.5 dpc, one group was sacrificed for tissue collection, the second group was sacrificed post-gestation, with additional samples collected post-MRI (n=6 per group). (B) Daily measurements of arterial pressure during 18 days of gestation in normal, preeclamptic, and treated preeclamptic mice. Statistical significance: *P<0.05, **P≤0.01, ***P≤0.001.

Preeclamptic mice exhibited a significant increase in arterial blood pressure starting from day 13.5 after plug and persisting until day 18.5. The highest peak was observed at day 15.5 reaching a mean arterial pressure of approximately 120 mmHg, which was significantly higher than the mean arterial pressure of 90 mmHg observed in normal gravid mice. Treatment with PC1 effectively prevented this increase and maintained blood pressure at normal levels (Figure 1B).

Urine samples were collected from each mouse on days 7.5 and 14.5 to assess proteinuria as an indicator of renal function. Total protein concentration and the albumin/creatinine ratio were determined. Preeclamptic mice showed significant hyperproteinuria compared to normal gravid mice, observed from the first to the third part of gestation (Figure S1). In terms of body weight preeclamptic and normal gravid mice exhibited a gradual increase throughout the gestation period, with a similar pattern of weight gain (Figure S1).

At day 18.5 of gestation one day prior to delivery, both the placenta and the corresponding fetuses were collected to assess placental efficiency. This parameter was decreased in PE mice compared to normal gravid mice (∼1.5-fold decrease), indicating impaired fetal growth associated with placental dysfunction. Treatment with the PC1 antagonist did not prevent the increase in proteinuria nor placental deficiency (Figure S1).

### PROKs levels in plasma and brain regions

Circulating levels of PROK1 and PROK2 were measured in mouse plasma at the end of gestation at day 18.5 and 8 months after gestation (Figure 2A and 2D). We observed an elevation of PROK1 and PROK2 concentrations in PE mice at E18.5. PROK1 levels reached 100 pg/mL whereas PROK2 levels were at 7.5 ng/mL. In comparison, normal pregnant mice exhibited lower levels of PROK1 (4 pg/mL) and PROK2 (3.8 ng/mL). Treatment with PC1 effectively prevented the elevation of PROK1 in preeclamptic mice and maintained their levels within the normal range of 4 pg/mL. Additionally, PC1 prevented the excessive increase of PROK2 in preeclamptic mice, reducing its levels to 5.8 ng/mL

**Figure 2:**
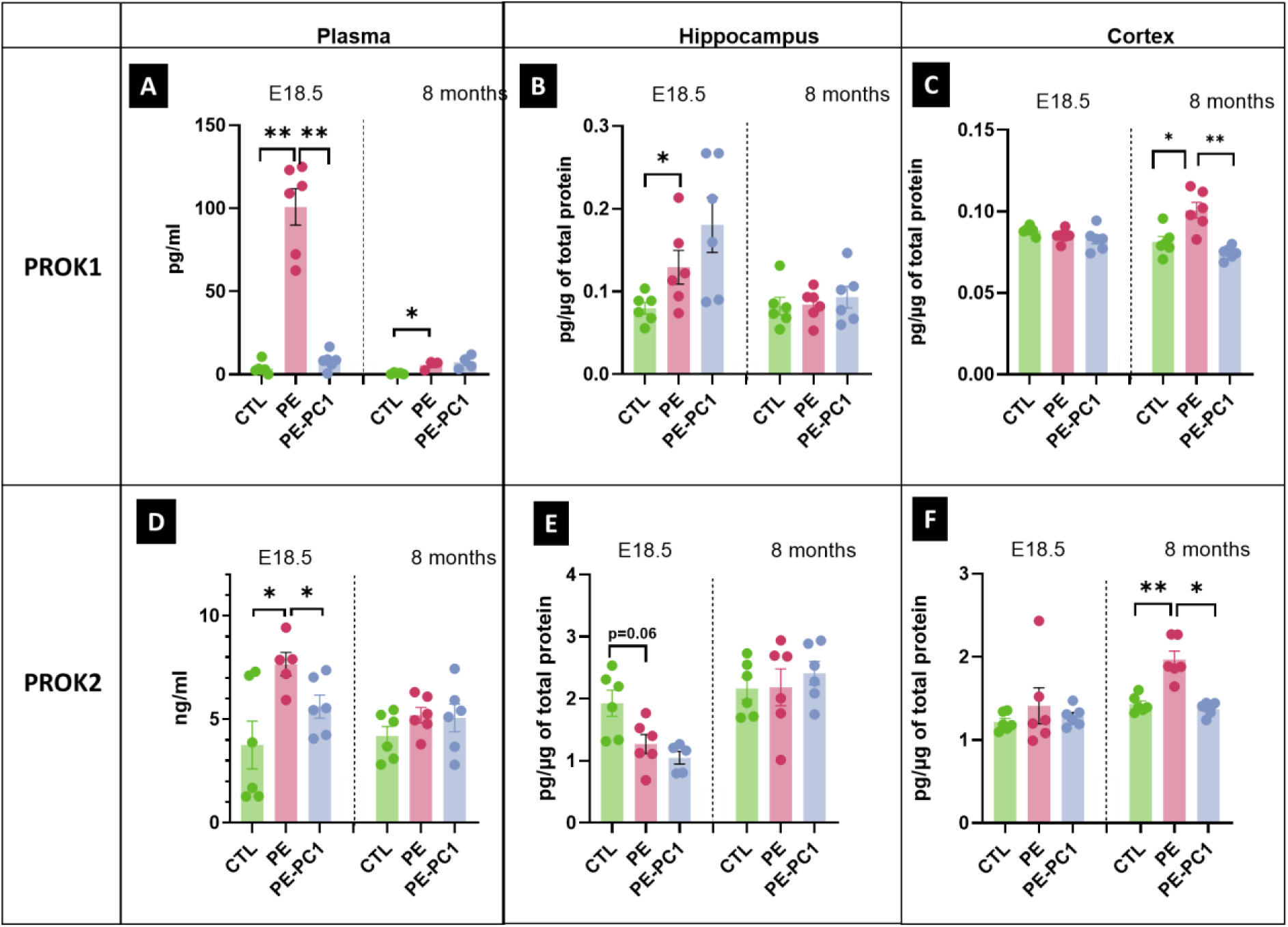
PROK1 and PROK2 concentrations in plasma (A,D), cortex (B,E) and hippocampus (C,F) from control mice (CTL), preeclamptic mice (PE) & preeclamptic mice treated with the antagonist of PROK-receptor (PE-PC1). Measurements were perfomed at E18.5 and 8 months postpartum. Data are presented as mean ± SEM (n=6 mice/group). Ordinary one-way ANOVA, corrected by Tukey’s test for multiple comparisons, was used for statistical analysis. Statistical significance is indicated as *P<0.05 and **P≤0.01.

In the long term, eight months after the preeclampsia phase, both PROK1 and PROK2 returned to near-normal levels in the plasma. PROK1 levels in preeclamptic mice decreased to approximately 10 pg/mL, but remaining significantly higher than the age-matched control mice levels of 4 pg/mL. PROK2 levels in PE mice decreased from 8 ng/mL to 4.5 ng/mL nearing the normal values of 4 ng/mL. Mice treated with PC1 maintained PROK1 and PROK2 levels within the normal range over the long term.

To determine the potential role of PROK1 and PROK2 during preeclampsia in cerebral changes, measurements of their concentrations were determined in various regions of the brain both during pregnancy and 8 months later (Figure 2B, C, E, F). At 18.5 days of preeclamptic gestation PROK1 and PROK2 were only modified in the hippocampus: PROK1 was significantly increased and PROK2 concentrations decreased (Figure 2B and 2E). In both cases, PC1 treatment did not prevent these changes. It should be noted that the concentrations of PROK2 in the control group of mice were 200 and 100 times higher than that of PROK1, respectively, in the hippocampus and the cortex. In the cortex no variation in the concentration of PROK1 and PROK2 was observed, as in the other regions examined (cerebellum, thalamus, and hypothalamus) (Figure 2C and 2F, and Figure S2). In all cases, the PROKR antagonist treatment had no effect on PROK levels (Figure 2 and S2).

In contrast, 8 months after the preeclampsia phase both PROK1 and PROK2 exhibited significantly elevated levels in the cortex compared to age-matched control mice (Figure 2C and 2F). The PC1 treatment applied during the preeclamptic gestation prevented the elevations in the concentrations for both PROK1 and PROK2, resulting in PROK1 and PROK2 levels in PE mice similar to those seen in age-matched control mice. In the hippocampus and the other examined areas no variation was observed at this time after preeclampsia (Figure 2B, 2E, and S2).

### Levels of expression of PROKs receptors in cortex and hippocampus

Western blots were performed to investigate PROKR expression in the cortex and hippocampus during the preeclampsia phase and 8 months later in the absence or presence of PC1 treatment (Figure 3). The PROKR1 blots showed three distinct immunoreactive bands at 37, 90, and 125 kDa (Figure S3). The most abundant PROKR1-immunoreactive band detected across the different brain tissues migrated at approximately 125 kDa, potentially representing a dimeric form of the receptor, as previously reported [39]. Therefore, we quantified both this band and the band detected at approximately 37 kDa, which corresponds to the predicted molecular weight of the native, unglycosylated PROKR1 receptor.

**Figure 3:**
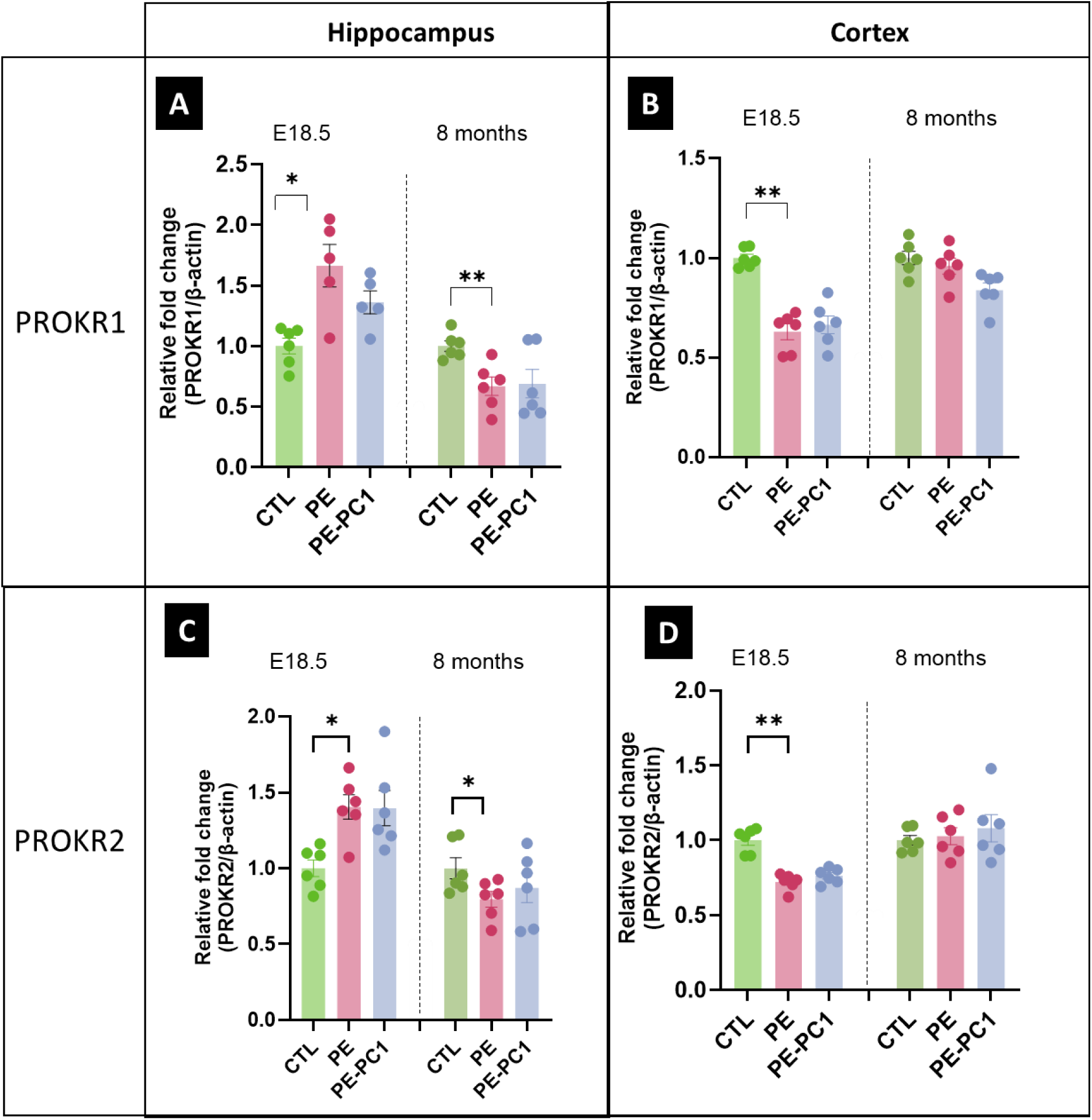
Expression of PROKR1 (A and B) and PROKR2 (C and D) in in the hippocampus (A and C) and the cortex (B and D) in control (CTL), preeclamptic mice (PE) and preeclamptic mice treated with anti PROK-receptor antagonist (PE-PC1). Measurements were performed at E18.5 and 8 months postpartum. Data are presented as PROKR1 and PROKR2 expression levels normalized to the corresponding β-actin signal, in respective brain regions (n=6 mice/group). Ordinary one-way ANOVA, corrected by Tukey’s test for multiple comparisons, was used for statistical analysis. Statistical significance is indicated as *P<0.05 and **P≤0.01.

PROKR2 blots displayed one band at 44 kDa. Expression of receptors was evaluated by the ratio intensity of PROKR1 or PROKR2 to actin bands

During the preeclampsia phase the intensity of the 125 kDa PROKR1-immunoreactive band and PROKR2 expression significantly increased in the hippocampus without any effect of PC1 (Figure 3A and 3C). Conversely, the expression of PROKR1 and PROKR2 significantly decreased in the cortex of preeclamptic mice, but without any significant preventive effect of PC1 treatment (Figure 3B and 3D).

Notably, in the hypothalamus of preeclamptic mice, a specific increase in PROKR2 appeared, which was prevented by the PC1 treatment (Figure S3).

Beyond 8 months after preeclampsia both PROKR1 and PROKR2 in the hippocampus showed significantly lower expression in preeclamptic mice than in normal mice, and in all cases, PC1 treatment had no effect compared to preeclamptic mice (Figure 3A and 3C). In the other regions it is worth noting that in the thalamus PROKR1 expression was specifically increased in preeclamptic mice compared to age-matched control mice and was significantly prevented by PC1 treatment (Figure S3). Moreover, in the hypothalamus, PROKR2 expression was significantly decreased in preeclamptic mice compared to normal mice, but PC1 treatment had no effect on their expression (Figure S3).

### MRI studies at 8 months post-gestation

To assess the long-term neurological consequences of preeclampsia we performed a high-resolution magnetic resonance imaging on animals 8 months postpartum using a 9.4 Tesla MRI scanner. This approach allowed the monitoring and characterization of cerebral lesions, including hemorrhagic and non-hemorrhagic abnormalities, assessed jointly on T2*-weighted and pre- and post-contrast T1-weighted images T2*-weighted imaging was employed to detect hemorrhagic lesions based on the paramagnetic properties of blood breakdown products. Specifically, T2* is highly sensitive to deoxyhemoglobin, methemoglobin, and hemosiderin which generate local magnetic field inhomogeneities that appear as hypointense regions on T2*-weighted images, often referred to as "black holes" [40]. Additionally, we used T1-weighted imaging with gadolinium (Gd-DOTA) contrast enhancement to evaluate blood-brain barrier (BBB) integrity. Since gadolinium-based contrast agents do not cross an intact BBB any signal enhancement in T1-weighted images following Gd-DOTA injection indicates localized BBB disruption reflecting increased vascular permeability in affected brain regions. Coronal brain MRI revealed focal abnormalities in preeclamptic mice (Figure. 4A). A well-demarcated lesion was observed in the caudoputamen, beneath the corpus callosum (Figure. 4A, row a, yellow arrows). This lesion was hyperintense on both pre- and post-contrast T1-weighted images, without corresponding signal loss on T2*-weighted images. T1 hyperintensity can reflect methemoglobin or non-hemorrhagic tissue constituents and is not specific for hemorrhage [41], the absence of corresponding T2* signal loss therefore leaves the nature of this lesion uncertain.

**Figure 4:**
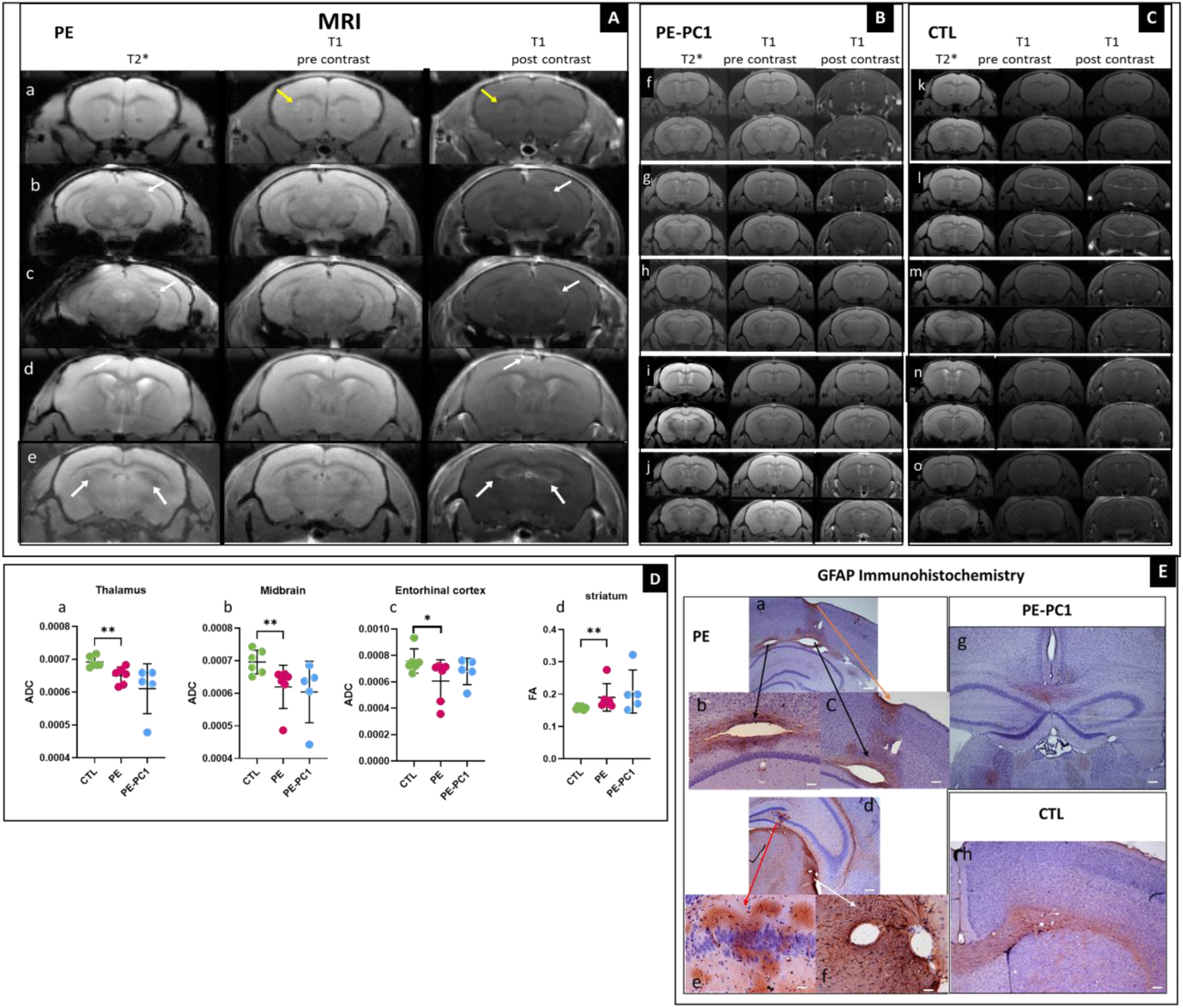
MRI Analysis. Representative of coronal brain T2*-weighted, pre- and post-contrast T1-weighted images obtained by MRI of 8 months postpartum from five preeclamptic mice (A, a-e), five PC1-treated mice (B, f-j) and five control mice (C, k-o) in multiple planes. (D) Statistical analyses of Apparent Diffusion Coefficient (ADC, a-c) and Fractional Anisotropy (FA, d) raw data from thalamus, midbrain, entorhinal cortex and striatum of PE, treated-PC1 and control mice. Statistical analyses were performed with Mann-Whitney test, p-value is considered significant for *p < 0.05, and ** p<0,01. *Immunohistochemistry*. (E) Immunohistochemistry and GFAP staining in PE mice ((a-f) revealing cortical lesion (orange arrow) and WM lesion (black arrows) associated with intense GFAP staining (e). Absence of lesions and weak GFAP staining are noticed in PE mice treated with PC1 (g) and in control mice (h). Images in panels a, d, g, and h were acquired at 2.5× magnification (scale bars, 200 µm), those in panels b and c at 5× magnification (scale bars, 100 µm), and those in panels e and f at 10× magnification (scale bars, 50 µm).

Additional abnormalities were detected in the hippocampus (Figure. 4A, rows b, c, and e, white arrows) and the parieto-temporal cortex (Figure. 4A, row d, white arrows). These regions exhibited focal hypointensities on T2*-weighted images, consistent with micro-hemorrhagic changes, together with increased signal intensity on post-contrast T1-weighted images (white arrows). Such T2* hypointensities in the hippocampus and parieto-temporal cortex are compatible with micro-hemorrhagic changes [40]. Increased signal on post-contrast T1 images, when confirmed relative to pre-contrast images and by subtraction, supports parenchymal enhancement and increased BBB permeability.

In contrast, no hemorrhagic lesions were observed in the corresponding images from age-matched control mice or PC1-treated preeclamptic mice (Figure. 4B, C). The images from PC1-treated mice appeared comparable to those from age-matched control mice.

Taken together, these findings support heterogeneous cerebral injury with a microhemorrhagic component and a distinct T1-hyperintense lesion potentially representing a nonhemorrhagic component [40,41].

Statistical raw data analyses were performed for every region from the atlas registered to the images. Significant differences between preeclamptic mice and control mice were observed for the apparent diffusion coefficient (ADC) and the fractional anisotropy (FA) (Figure 4D). ADC analysis from the different regions revealed a significant decrease only in the thalamus, midbrain, and entorhinal cortex indicating an alteration of water circulation that might reveal the presence of lesions in these regions (Figure 4D-a, b, c). Statistical FA raw data analyses showed increased values only in the striatum from PE animals, consistent with altered tissue microstructure in this region (Figure 4D-d). PC1 treatment did not allow recovery from PE perturbations in these regions for ADC and FA analyses

To assess whether inflammation coexist with lesions observed by MRI, immunostaining using glial fibrillary acidic protein (GFAP) was performed on perfused brains (Figure 4E). GFAP is an intracellular intermediate filament protein expressed by astrocytes in the central nervous system and serves as a marker for visualizing reactive astrocytes and estimating the degree of brain injury [42]. Immunostaining of the brains of PE mice showed GFAP immunostaining associated with large lesions in the cortex (orange arrow) and to white matter lesions (WML) (Figure 4E-a, b, c; black arrows). A strong GFAP signal was also observed in the hippocampus at the level of the CA4 and the gyrus dentate (Figure 4E-d, e; red arrow) and in the lateral thalamic tissue close to the lateral ventricle (Figure 4E-d, f; white arrow). Compared to control mice, no such GFAP immunostaining was evidenced (Figure 4E-g, h), supporting an association between experimental preeclampsia and persistent astrocyte reactivity, long term after the acute phase of the pathology. Notably, these results showed that treatment with the antagonist of the prokineticin receptor, PC1, prevented lesions and associated astrocyte reactivity.

## 4. Discussion

### Long-term neurological consequences of preeclampsia

Preeclampsia is increasingly recognized as a multisystem disorder with effects that extend beyond pregnancy, particularly in the central nervous system (CNS). Our study demonstrates persistent cerebral abnormalities in the STOX1A model (i.e; where phenotypes develop in WT females and are exclusively caused by the placenta-fetal overexpression of STOX1A), with MRI findings consistent with BBB disruption and astrocyte reactivity. These findings align with clinical observations of increased risks of stroke, white matter lesions (WML), and cognitive decline in women with a history of PE [11, 12, 43,44]. The hippocampal and parieto-temporal T2* hypointensities are compatible with micro-hemorrhagic injury, whereas the caudoputamen lesion, hyperintense on T1 without corresponding T2* signal loss, may represent a non-hemorrhagic component [0,41]. Intrinsic pre-contrast T1 hyperintensity was distinguished from post-Gd-DOTA enhancement supporting increased BBB permeability when localized to the parenchyma. The coexistence of these findings at 8 months postpartum supports persistent cerebrovascular injury beyond the acute hypertensive phase. This distinction is relevant because PE is associated with an increased risk of ischemic stroke during pregnancy and the postpartum period, and a more modest elevation of long-term risk [43, 45,46].

### The prokineticin system emerged as a key mediator of preeclampsia associated cerebral alterations

During PE, we observed a 25-fold increase in circulating PROK1 and a 2-fold increase in PROK2 levels which were normalized by PC1 treatment. This supports previous findings of elevated PROK1 in PE [20,21] and is consistent with recent reports that PROK2 is also dysregulated in PE [47]. The normalization of prokineticin levels following PC1 treatment suggests that PROKR1 for which PC1 has an approximately 30-fold higher affinity than for PROKR2 [38, 48], may contribute to the hypertensive and vascular alterations observed in PE, although the relative contributions of PROKR1 and PROKR2 cannot be established from PC1 treatment alone. This is consistent with studies showing that PROK1 and PROK2 can increase BBB permeability *in vitro* [49], and that the antagonists PC7 and PKR-A attenuate this effect in that model. Thus, taken together the present findings support a possible link between prokineticin signaling and PE-associated cerebrovascular alterations.

### Mechanisms underlying the protective effects of PC1 on cerebral lesions

Treatment with PC1 prevented the development of focal brain lesions in mice with pre-eclampsia, as well as the appearance of abnormalities detectable by MRI, and, concomitantly, the increase of inflammation, revealed by astrocyte activation, where lesions were predominantly localized in PE mice. Astrogliosis is associated with neurological damage, cognitive decline and dementia [13, 38]. The protective effects of PC1 may involve a reduction in PROK-mediated disruption of the BBB and neuroinflammation [49], although its effects on tissue microstructure remain incompletely understood. However, the contribution of direct central effects of PC1 cannot be separated from indirect effects resulting from normalization of gestational blood pressure.

The differential effects of PC1 across brain regions highlight the complexity of prokineticin signaling. PC1 prevented the long-term elevation of PROK1 and PROK2 in the cortex but did not reverse the decrease in PROKR1 and PROKR2 expression in the hippocampus, 8 months postpartum. Such regional responses may reflect differences in receptor signaling or local tissue conditions. Divergent actions of PROKR1 and PROKR2 have also been described in cardiac physiology and pathology [50], supporting the importance of tissue context without establishing an equivalent mechanism in the brain. PROKR2 can form dimers [39], and its activation may trigger different downstream effects according to cellular context. PROK2 has been shown to promote neuroprotection in Parkinson’s disease models and an astrocyte response described as A2-like [31, 51], whereas PROK1 may contribute to BBB disruption [49]. Regarding the molecular identity of PROKR1 implicated, subcellular localization, and functional significance of the approximately 125 kDa PROKR1-immunoreactive band remain to be established and warrant further investigation using independent validation approaches. Further studies should clarify how these regional expression patterns relate to receptor activity and the neurological effects in a PE setting.

### Correlation between MRI findings and inflammatory markers

GFAP immunostaining revealed reactive astrocytosis in brain regions affected by PE. Its concordance with MRI abnormalities supports an associated neuroinflammatory component, without establishing the cause of each MRI signal change. In human studies, WML are commonly observed after PE, particularly in the frontal and temporal lobes [11, 12], and are associated with cognitive impairment and dementia [4–6, 9, 13]. These observations suggest that vascular injury and neuroinflammation may contribute to both experimental PE-associated brain injury and long-term neurological complications in women.

The ADC and FA alterations provide complementary evidence of microstructural changes. Decreased ADC in the thalamus, midbrain, and entorhinal cortex suggests restricted water diffusion that may reflect cellular swelling or tissue damage [52]. Increased FA in the striatum indicates altered tissue microstructure [52,53]. Their persistence despite PC1 treatment suggests that prokineticin antagonism alone does not fully prevent these changes. Other pathways, including oxidative stress and endothelial dysfunction, may contribute [54]. Investigating their interaction with prokineticin signaling could help explain the residual abnormalities and identify complementary therapeutic targets.

### Extrapolation to human studies and clinical implications

The findings of this study support the hypothesis that persistent cerebrovascular alterations contribute to the long-term neurological consequences of PE. Cerebral abnormalities consistent with BBB disruption remained detectable at 8 months postpartum in mice, suggesting that vascular barrier alterations may persist beyond the hypertensive pregnancy. The regional dysregulation of prokineticins, together with their reported effects on BBB integrity [49], provides a possible mechanistic link between PE and this persistent vascular vulnerability. This interpretation is consistent with the long-lasting hippocampal vascular dysfunction and memory impairment reported in a rat model of PE [32], as well as with white matter abnormalities and increased stroke risk in women with a history of PE [11, 12, 43,45,46]. Cohort studies and a meta-analysis further support an association between PE and later vascular dementia, whereas the association with Alzheimer’s disease is less consistent [6, 7, 9]. Our principal hypothesis is therefore that persistent vascular injury after PE may contribute to cerebral small vessel damage and vascular cognitive impairment, and may represent a potential substrate for later vascular cognitive impairment [10]. The residual diffusion abnormalities despite PC1 treatment may reflect lasting tissue injury, although they are not specific to a vascular mechanism. Longitudinal assessments of cerebrovascular function and cognition are needed to test this hypothesis, as dementia was not assessed.

In the present study, PC1 treatment was initiated on gestational day 12, immediately before the rise in blood pressure observed on day 13, and prevented both gestational hypertension and the long-term MRI-detected lesions and associated astrogliosis. This timing precludes distinguishing direct cerebrovascular effects of PROKR antagonism from protection secondary to blood pressure control. Both mechanisms may contribute to the observed benefits. Indeed, STOX1A overexpression disrupts placental mitochondrial function and redox homeostasis, impairing adaptation to hypoxia and favoring oxidative and nitrosative stress [34,35]. These disturbances may promote the release of inflammatory and antiangiogenic mediators into the maternal circulation, compromising endothelial NO signaling and enhancing vasoconstrictor responses, thereby increasing vascular resistance and sustaining hypertension [33,55,56]. In this context, PC1, a non-peptide prokineticin receptor antagonist [37], could attenuate prokineticin-mediated vascular dysfunction while also protecting blood–brain barrier (BBB) integrity. Supporting the latter possibility, our previous work demonstrated that PROKR antagonists prevented PROK1- and PROK2-induced increases in permeability in an *in vitro* BBB model [49]. PC1 may therefore combine systemic haemodynamic benefits with direct antagonism of prokineticin signaling at the cerebrovascular endothelium, although the respective contributions of these mechanisms remain to be determined.

In summary, this study demonstrates persistent cerebral abnormalities in the STOX1A model following preeclampsia, including MRI features compatible with cerebrovascular injury, BBB permeability alterations, and regional microstructural changes, accompanied by persistent astrocyte reactivity. Gestational PC1 treatment attenuated selected long-term cerebral abnormalities and prevented the long-term cortical elevation of PROK1 and PROK2. These findings support a potential association between prokineticin signaling and persistent PE-associated cerebrovascular alterations. The present study focused on structural and molecular changes, however, further behavior and cognitive assessments are needed to determine their consequences and whether PC1 confers lasting neurological benefits. Thus, this study highlights our understanding of the neurological impact of PE, and the therapeutic potential of an intervention targeting the prokineticin system.

## Acknowledgments

We would like to express our sincere gratitude to BAMA-TOUPET Soumalamaya and MAGALLON Charlène, the skilled zootechnicians, for their dedicated work and expert care of the animals at the animal facility. Their technical support and commitment were essential for the successful completion of this study. We also extend our thanks to Nora COLOMB and Camille BEETS from the IRMaGe MRI platform for their expertise and assistance in performing the high-resolution MRI acquisitions. Their contributions were invaluable for the neurological assessments presented in this work.

This study was supported by funding from The University of Grenoble Alpes for HY scholarship, from the CERCOG@UGA (ANR-15-IDEX-0002), and the Grenoble Alliance for Integrated Structural & Cell Biology, a program from the Chemistry Biology Health, Graduate School of University Grenoble Alpes for IK scholarship and supports (ANR-17-EURE-0003).

## Authors contribution

HY: Conceptualization, experimental design, experiments, data analysis, data interpretation, data curation, original draft-FS: experimental design, experiments, technical assistance-IK: experiments, data analysis, data interpretation, data curation, technical assistance. ML: data analysis, data interpretation, data curation. OM: experimental design, experiments, data analysis, data curation, technical assistance. DV: Conceptualization. MB: Conceptualization, data interpretation. NA: Conceptualization, experimental design, data interpretation. CM: Conceptualization, experimental design, experiments, data analysis, data interpretation, data curation, original draft, revised draft and editing.

The authors declare that they have no competing interests.

## Abbreviations

PE: preeclampsia
PROK1 and PROK2: prokineticin 1 and prokineticin 2
PROKR1 and PROKR2: prokineticin receptor 1 and prokineticin receptor 2
CTL: control
BBB: blood brain barrier
ADC: apparent diffusion coefficient
FA: fractional anisotropy

## Supplementary results

**Figure S1.**
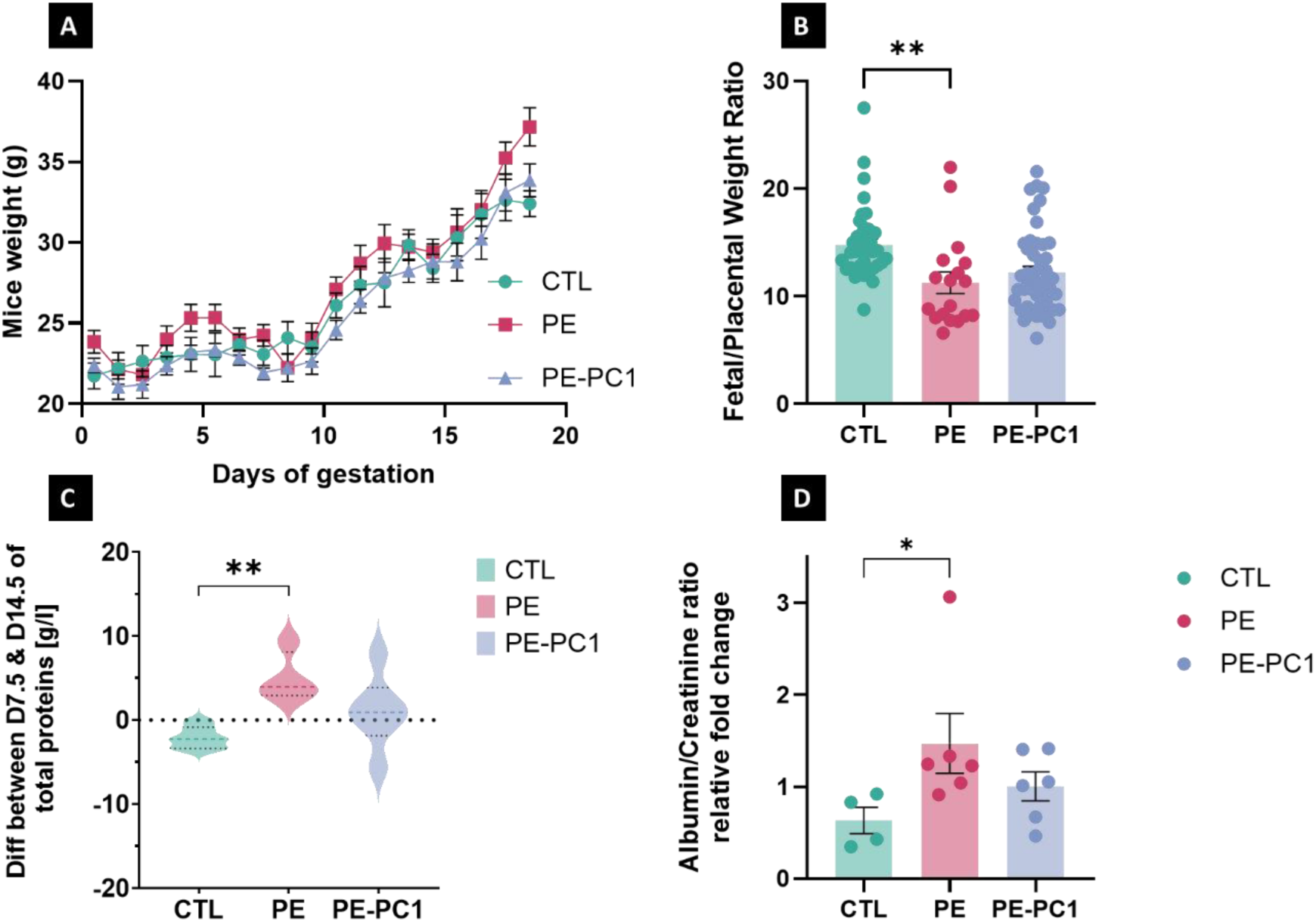
Validation of preeclamptic STOX1 model : Control, preeclamptic and PC1-treated preeclamptic mice were followed during gestation, for body weight (A), fœtal efficency (B), protenuria (C) and albumin-creatinine ratio (D). Ordinary one-way ANOVA, corrected by Tukey’s test for multiple comparisons, was used for statistical analysis. Statistical significance is indicated as *P<0.05 and **P≤0.01.

**Figure S2:**
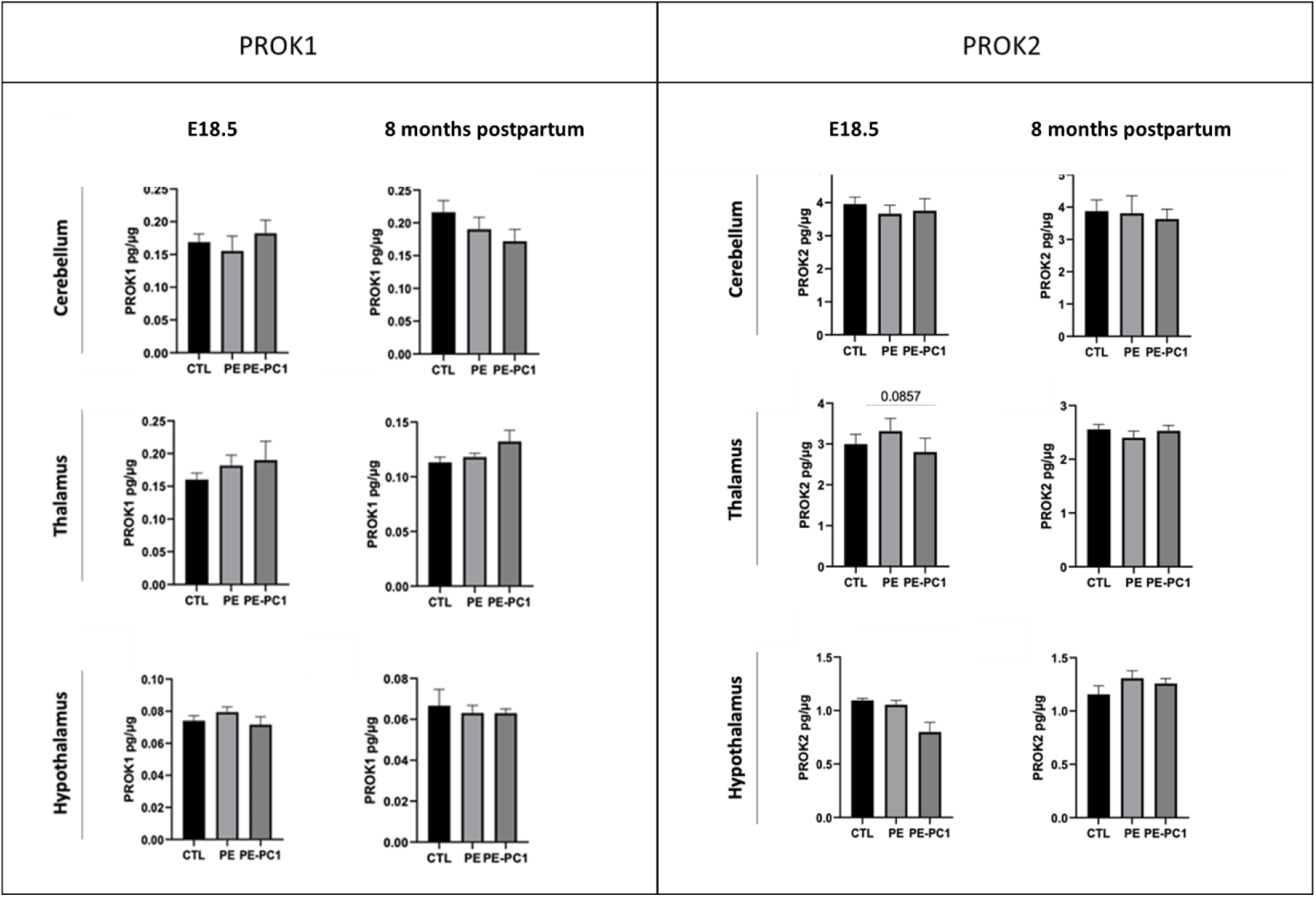
PROK1 and PROK2 concentration levels in brain regions. Concentration of PROK1 and PROK2 in the cerebellum, thalamus and hypothalamus of preeclamptic (PE), preeclamptic treated-PC1 (PE-PC1), and control (CTL) mice, 1 day prepartum and 8 months postpartum. Data are expressed as mean ± SEM of the amount of PROK1 or PROK2 in picograms per microgram of total protein in the respective brain regions (n=6 mice/group). Statistical analysis was performed using ordinary one-way ANOVA corrected by Tukey’s test for multiple comparisons and statistical significance is denoted as *P<0.05, **P≤0.01, ***P≤0.001.

**Figure S3 :**
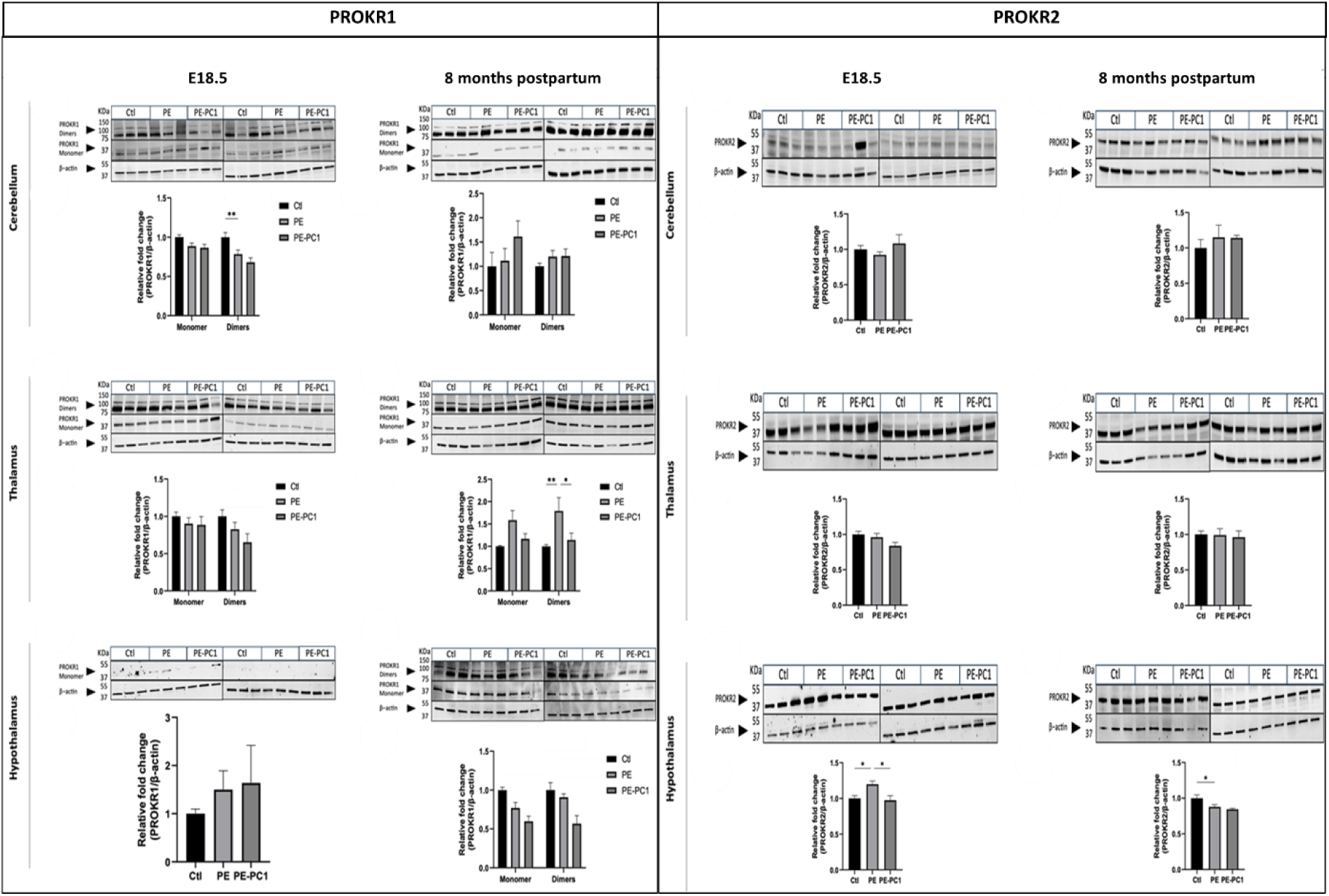
Expression of PROKR1 and PROKR2 in cerebellum, thalamus and hypothalamus of preeclamptic (PE), treated, and control (CTL) mice. Western blot images are shown along with their respective quantifications for PROKR1 monomer and oligomers, and PROKR2 expression. PROKR1 and PROKR2 expression is examined at 18.5 days of gestation and after 8 months of gestation in the corresponding brain regions. Data are expressed as mean―SEM (n=6 mice/group). The statistical comparison was performed using a two-way ANOVA test corrected by Tukey test for multiple comparisons, and statistical significance is indicated as *P<0.05, **P≤0.01, ***P≤0.001.

**Figure S4.**
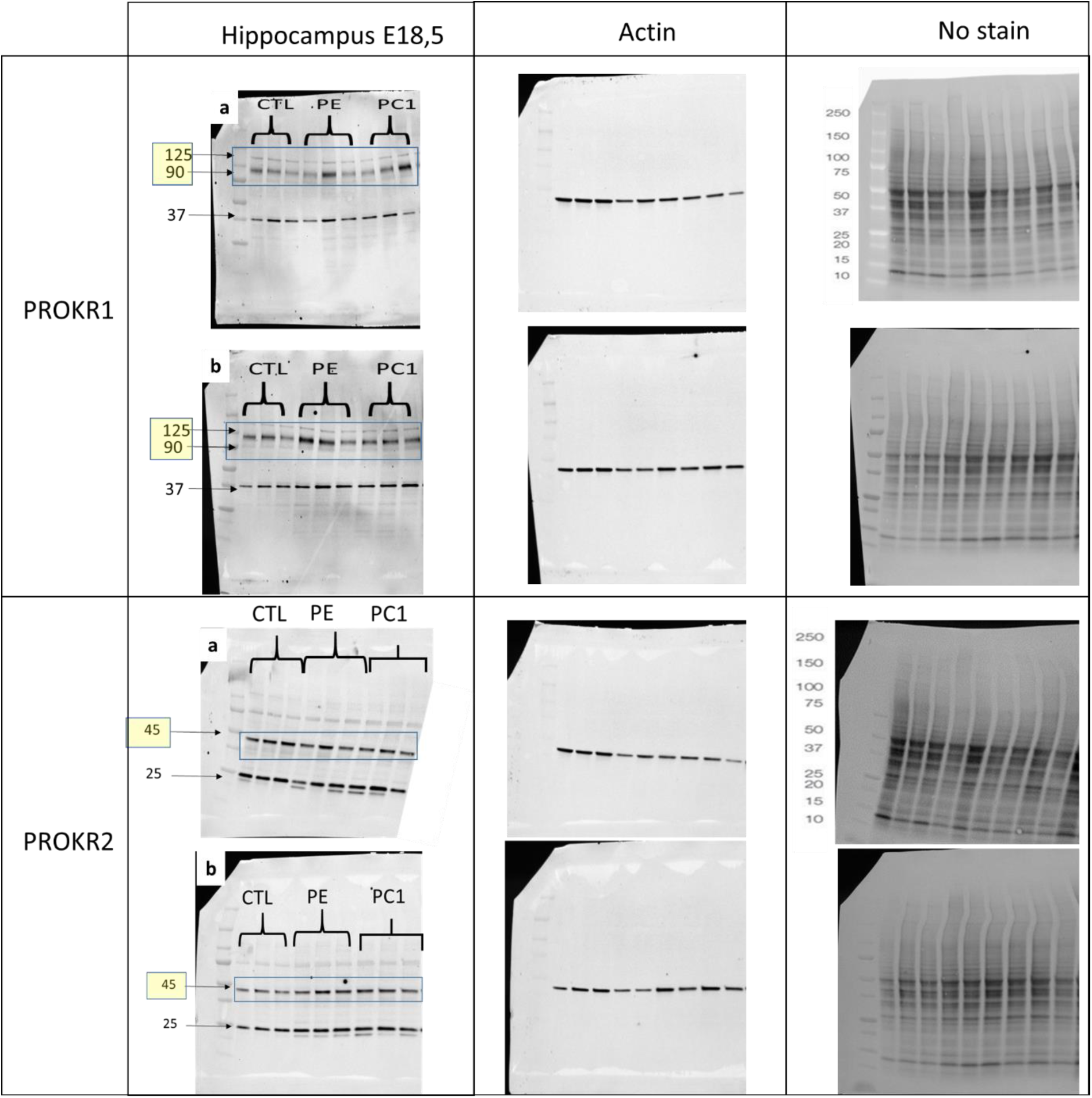
Complete Western blot membranes for PROKR1 and PROKR2 in the maternal hippocampus at E18,5. Full membrane images show PROKR1 (upper panels) and PROKR2 (lower panels), together with the corresponding β-actin loading controls and total-protein images. Membranes a and b each contain samples from three age-matched control pregnant mice (CTL), three preeclamptic mice (PE), and three PC1-treated preeclamptic mice (PC1), giving six individual animals per group. Animal identifiers are provided in the accompanying table. PROKR1-immunoreactive bands were detected at approximately 37, 90, and 125 kDa; the 125-kDa band was quantified. The PROKR2 band selected for quantification migrated at approximately 45 kDa. Molecular weights are indicated in kDa. Protein expression was normalized to the corresponding β-actin signal.

**Figure S5.**
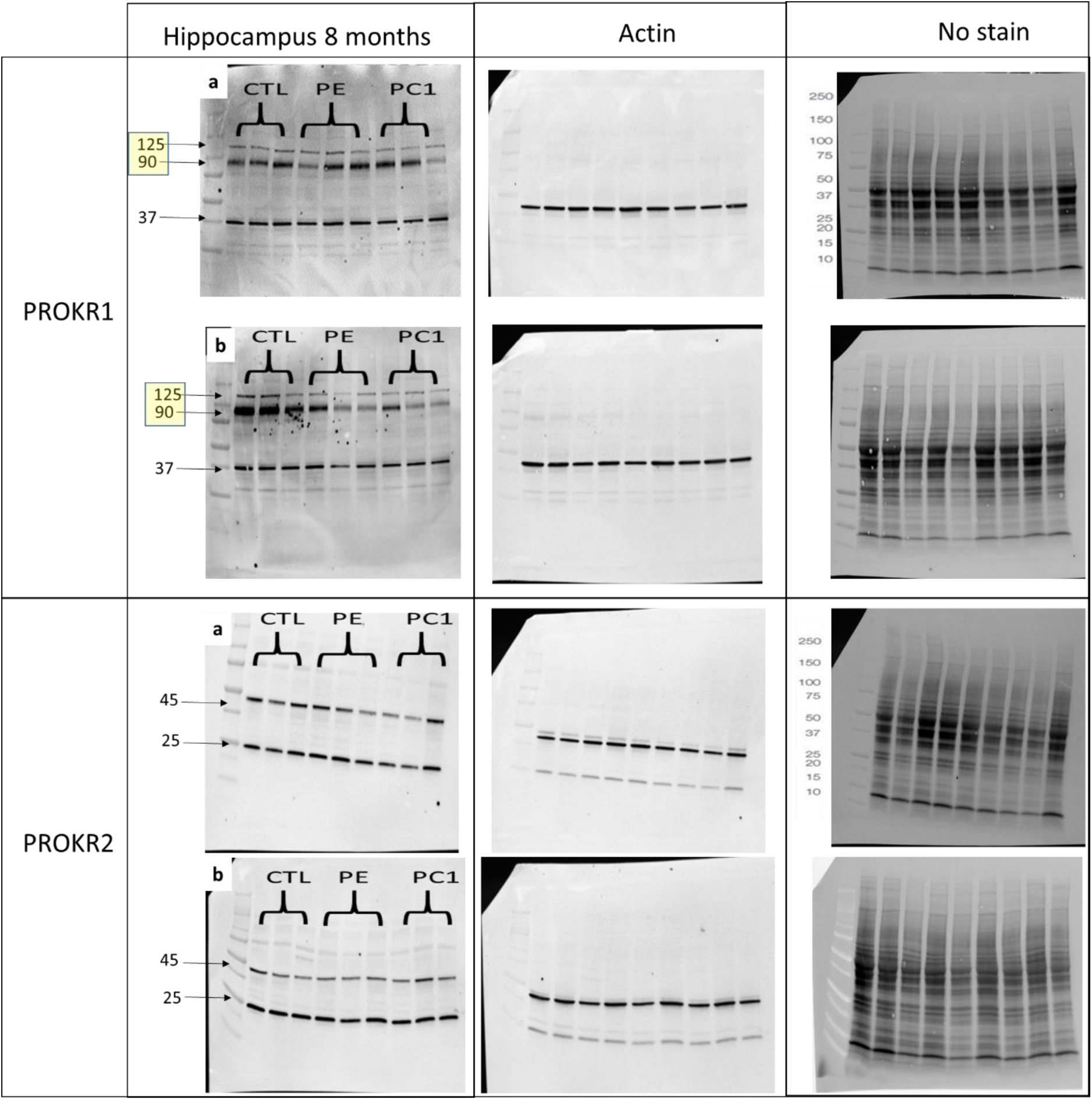
Complete Western blot membranes for PROKR1 and PROKR2 in the maternal hippocampus at 7 months postpartum. Full membrane images show PROKR1 (upper panels) and PROKR2 (lower panels), together with the corresponding β-actin loading controls and total-protein images. Membranes a and b each contain samples from three age-matched control pregnant mice (CTL), three preeclamptic mice (PE), and three PC1-treated preeclamptic mice (PC1), giving six individual animals per group. Animal identifiers are provided in the accompanying table. PROKR1-immunoreactive bands were detected at approximately 37, 90, and 125 kDa; the 125-kDa band was quantified. The PROKR2 band selected for quantification migrated at approximately 45 kDa. Molecular weights are indicated in kDa. Protein expression was normalized to the corresponding β-actin signal.

**Figure S6.**
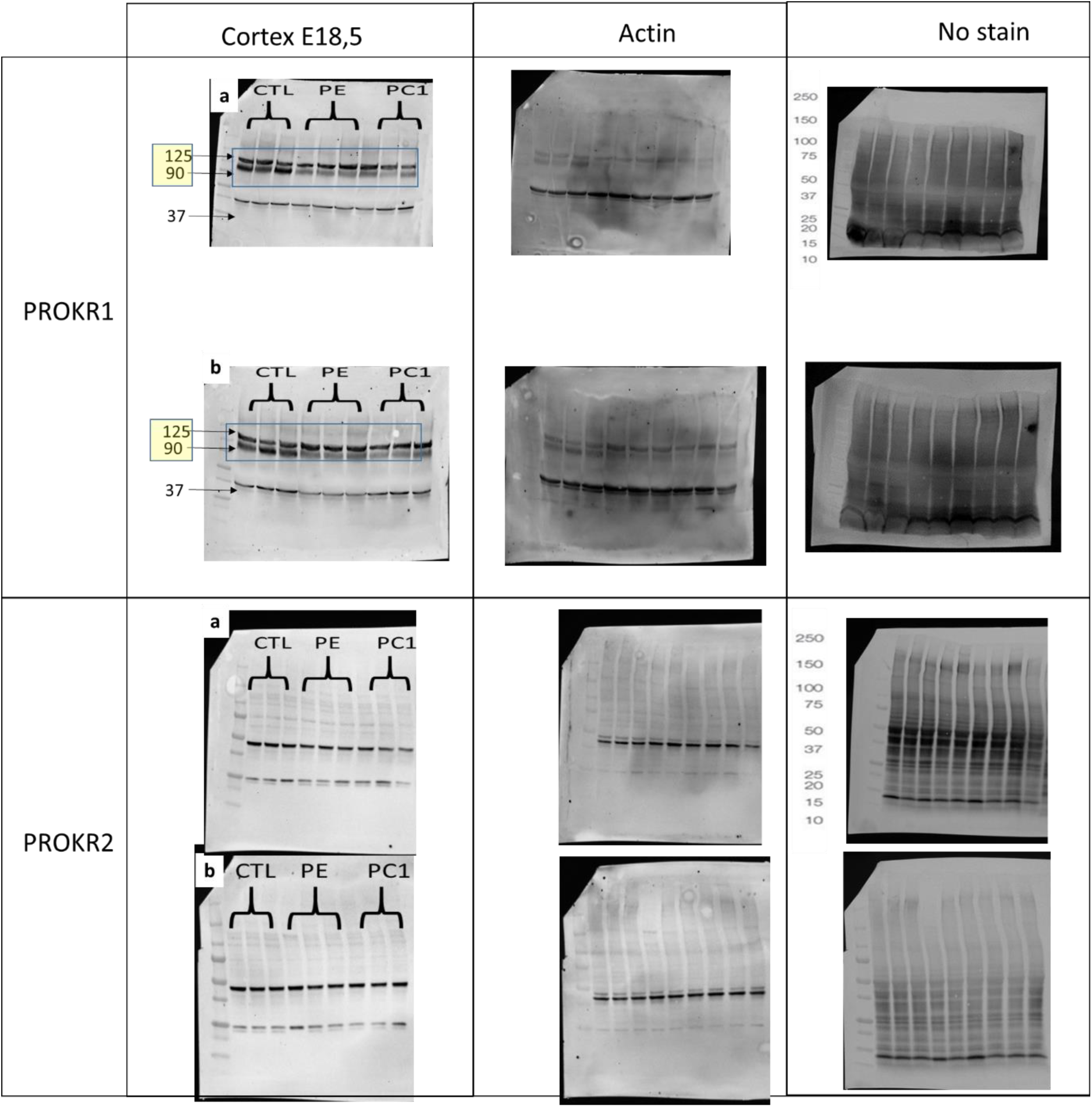
Complete Western blot membranes for PROKR1 and PROKR2 in the maternal cortex at E18,5. Full membrane images show PROKR1 (upper panels) and PROKR2 (lower panels), together with the corresponding β-actin loading controls and total-protein images. Membranes a and b each contain samples from three age-matched control mice (CTL), three preeclamptic mice (PE), and three PC1-treated preeclamptic mice (PC1), giving six individual animals per group. Animal identifiers are provided in the accompanying table. PROKR1-immunoreactive bands were detected at approximately 37, 90, and 125 kDa; the 125-kDa band was quantified. The PROKR2 band selected for quantification migrated at approximately 45 kDa. Molecular weights are indicated in kDa. Protein expression was normalized to the corresponding β-actin signal.

**Figure S7.**
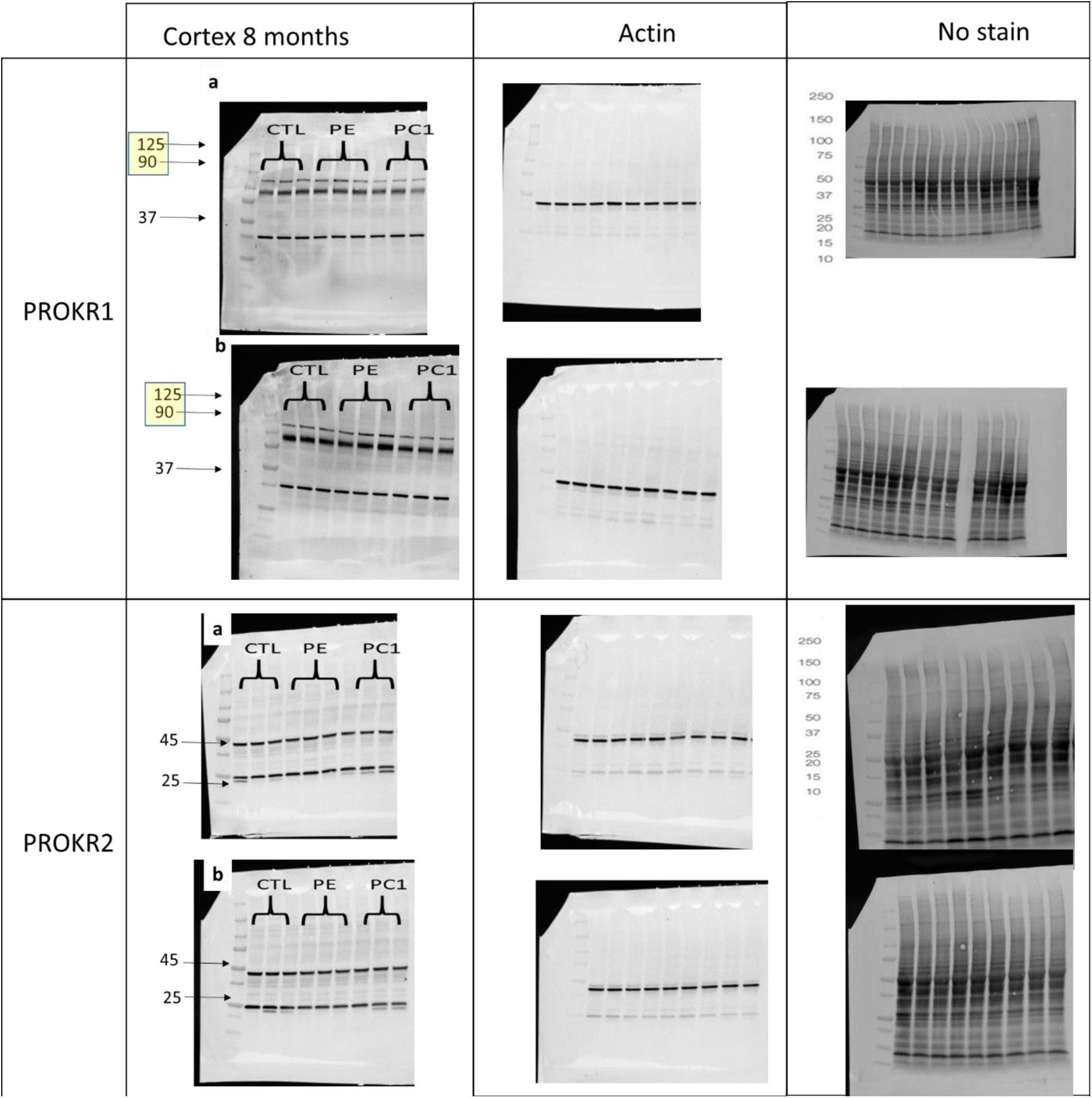
Complete Western blot membranes for PROKR1 and PROKR2 in the maternal cortex at 7 months postpartum. Full membrane images show PROKR1 (upper panels) and PROKR2 (lower panels), together with the corresponding β-actin loading controls and total-protein images. Membranes a and b each contain samples from three age-matched control pregnant mice (CTL), three preeclamptic mice (PE), and three PC1-treated preeclamptic mice (PC1), giving six individual animals per group. Animal identifiers are provided in the accompanying table. PROKR1-immunoreactive bands were detected at approximately 37, 90, and 125 kDa; the 125-kDa band was quantified. The PROKR2 band selected for quantification migrated at approximately 45 kDa. Molecular weights are indicated in kDa. Protein expression was normalized to the corresponding β-actin signal.

